# Mechanistic Insights into Na^+^-dependent HCO_3_^−^ Transport by NBCn2 (SLC4A10)

**DOI:** 10.64898/2026.03.09.710040

**Authors:** Lasse M. Desdorf, Amanda D. Stange, Helle H. Damkier, Birgit Schiøtt, Jeppe Praetorius, Anna L. Duncan

## Abstract

The 3D structure and mechanism of action are unknown for the integral plasma membrane transport protein Solute Carrier 4A10, which has been characterized functionally as an electroneutral Na^+^:HCO_3_^−^ cotransporter. We used structure prediction and molecular dynamics simulations to study the binding of the transported ions to the Solute Carrier 4A10 protein and suggest a model of sequential binding of Na^+^ followed by HCO_3_^−^ to the ion binding domain. The binding of HCO_3_^−^ to the protein appears to depend absolutely on Na^+^ binding. Conversely, binding of HCO_3_^−^ stabilizes the interaction between Na^+^ and its binding site. This allows the subsequent conformational changes of the Solute Carrier 4A10 protein and, thus, ion translocation. Measurements of intracellular pH and Na^+^ concentration revealed the dependence of Na^+^ on HCO_3_^−^ transport. The study lays the necessary foundation for advanced analysis of ion translocation and the development of selective transport inhibitors of Solute Carrier 4A10 and other proteins of the protein family of HCO_3_^−^ transporters.

## INTRODUCTION

Selective net transport of HCO_3_^−^ across cell membranes is governed by integral proteins belonging to the Solute Carrier (SLC) 4A family in mammals and beyond in the animal kingdom. The human SLC4A10 (NBCn2) is an electroneutral co-transporter of Na^+^ and HCO_3_^−^ into cells driven by the chemical Na^+^ gradient. NBCn2 is expressed in the brain and epithelial cells of the kidney and choroid plexus, where it is crucially involved in regulating both the intracellular and local extracellular environments, secretion, and absorption^1-3^. Thus, human loss-of-function mutations in NBCn2 lead to severe diseases with neurodevelopmental disorders, small brain ventricles, autism, and seizures^4,5^. By contrast, NBCn2 hyperactivity is implicated in conditions such as stroke, hydrocephalus, and trauma^6,7^. The need for efficient clinical control of NBCn2 activity is evident, but hopes for future therapies are hampered by the lack of deep knowledge of NBCn2 structure, ion binding, and transport dynamics.

NBCn2 is natively expressed as a homodimer and has two chains, A and B (Figure 1). It is a transmembrane protein consisting of 14 transmembrane helices, an extracellular loop region, an N-terminal region, and a C-terminal region. In these simulations, the N- and C-terminal regions have been truncated and will not be discussed further, due to the lack of structural data and relevance to the binding region of NBCn2. Currently, no structures have been solved of NBCn2, however, multiple structures exist of other SLC4A family members. As of writing this article, cryo-EM structures have been solved for the SLC4A1, A2, A3, A4, A8, and A11 in various conformational states^8-15^. These lay the foundation for structure predictions and computational modelling of the NBCn2 homodimer. It is proposed that the SLC4A family functions using the elevator-type mechanism, similar to other transmembrane homomers^16-20^.

**Figure 1.**
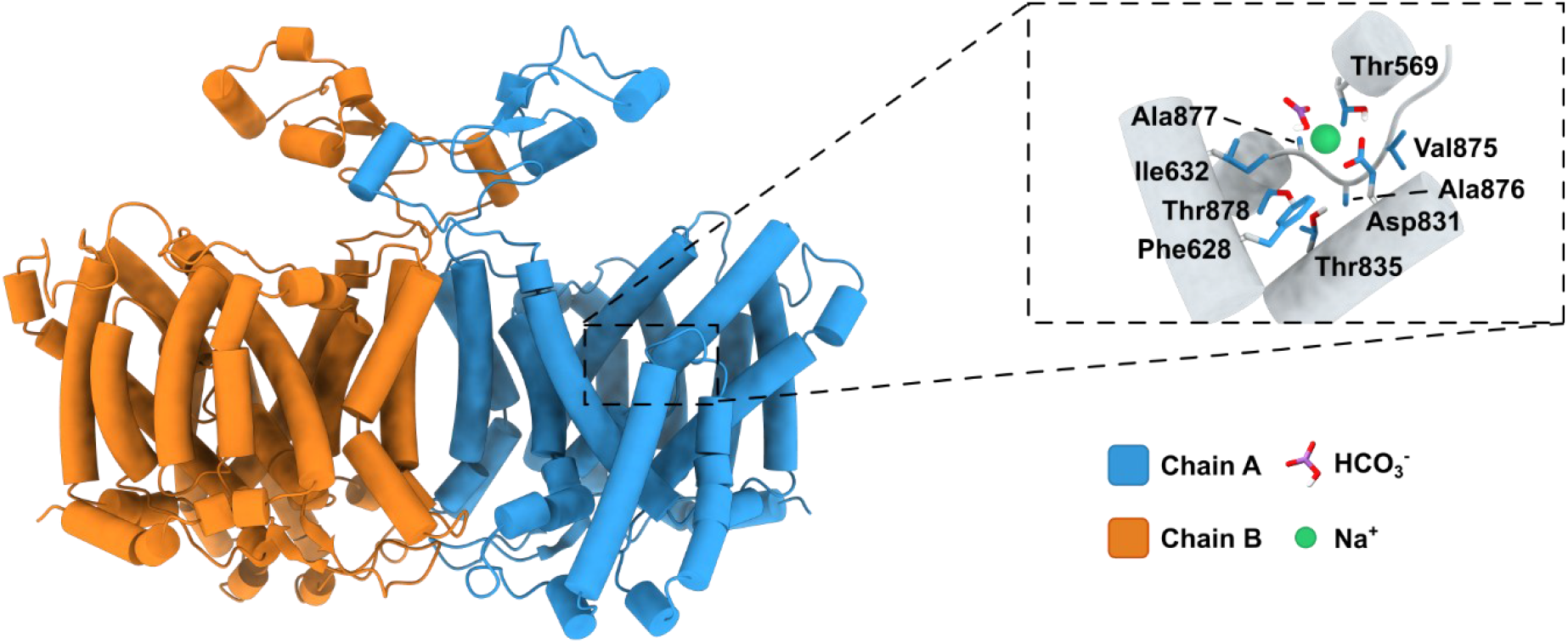
Structure of the NBCn2 homodimer. To the left, the NBCn2 homodimer (SLC4A10 protein) predicted by AlphaFold2. Chain A in blue and chain B in orange. The inset shows the magnified binding site for HCO_3_^−^ (purple) and Na^+^ (green) among key binding site residues that coordinate them.

It is currently unknown how NBCn2 binds and transports its substrate ions, HCO_3_^−^ and Na^+^. Additionally, the stoichiometry has not been fully resolved, and key residues for binding and stabilizing the ions have not been revealed. In this study, we aim to uncover the substrate dependence in NBCn2 and the ion stability by identifying the key binding site residues involved. In addition, we aim to correlate the key binding site residues in NBCn2 with other SLC4A family members to obtain information about the subfamily member nuances and dependence on Na^+^.

We utilize molecular dynamics (MD) simulations to study the stability and binding of the substrate ions of NBCn2. We simulated NBCn2 as a homodimer in three different system setups that contain either: HCO_3_^−^ and Na^+^; only HCO_3_^−^; or only Na^+^ in the binding site. Our MD simulations highlight the necessity of Na^+^ in the binding site to stabilize HCO_3_^−^. Removing Na^+^ destabilizes HCO_3_^−^ and leads to its rapid unbinding. Removing HCO_3_^−^ affects the stability of Na^+^, although, to a lesser extent, which is likely caused by Na^+^ being bound deeper in the binding site. We aligned the sequences of the SLC4A family to compare the conservation of key binding site residues identified by our simulations. Key binding site residues that interact with HCO_3_^−^ are conserved throughout, while Na^+^ interacting residues are only conserved in Na^+^-dependent family members. These results correlate with previous experimental findings, which show the inability to transport HCO_3_^−^ in Na^+^-depleted environments. We propose a sequential binding mechanism, where Na^+^ is bound first and then HCO_3_^−^ can bind to the outward facing binding domain. NBCn2 is then able to undergo a conformational change and release the ions into the cell’s cytoplasm from the inward facing domain of the protein.

## RESULTS

We generated a single starting structure of NBCn2 without substrate using Colabfold^21^. The results are based on MD simulations of three different system setups based on the substrate placed in the binding site of the starting structure. The systems are named: dual-substrate, HCO_3_^−^-only, and Na^+^-only (Table 1).

**Table 1.** Naming of system setups.

| System name | Ion in the binding site |  |
| --- | --- | --- |
| | $\text{HCO}_3^-$ | $\text{Na}^+$ |
| Dual-substrate | Yes | Yes |
| $\text{HCO}_3^-$-only | Yes | No |
| $\text{Na}^+$-only | No | Yes |

### HCO_3_ ^−^ binding requires the presence of Na^+^

We simulated two different systems containing HCO_3_^−^ (Table 1) to assess the importance of Na^+^ on the HCO_3_^−^ stability and binding. The stability of HCO_3_^−^ was investigated by calculating the distance between the binding site and HCO_3_^−^ over the entire simulation trajectory. The binding of HCO_3_^−^ was analyzed by calculating the protein–ion fingerprint, and key residues were defined as residues that interacted more than 5% of the total simulation time.

HCO_3_^−^ was found to be stable in the binding site when Na^+^ was bound (dual-substrate system) (Figure 2A). This was found across all repeats, independent of which monomeric chain it was located in. HCO_3_^−^ did not have a distance value above 1.0 nm, which means that it stayed in the original position during the simulation (Figure 2A, magnified insert). Removing Na^+^ from the binding site (the HCO_3_^−^-only system) notably reduced the stability of HCO_3_^−^ (Figure 2B). HCO_3_^−^ unbinds before 100 ns, as indicated by the sharp increase in distance, to a value indicative of HCO_3_^−^ entering the bulk phase. This was observed across repeats and monomeric chains. This indicates that the presence of Na^+^ in the binding site is crucial for the stability of HCO_3_^−^.

**Figure 2.**
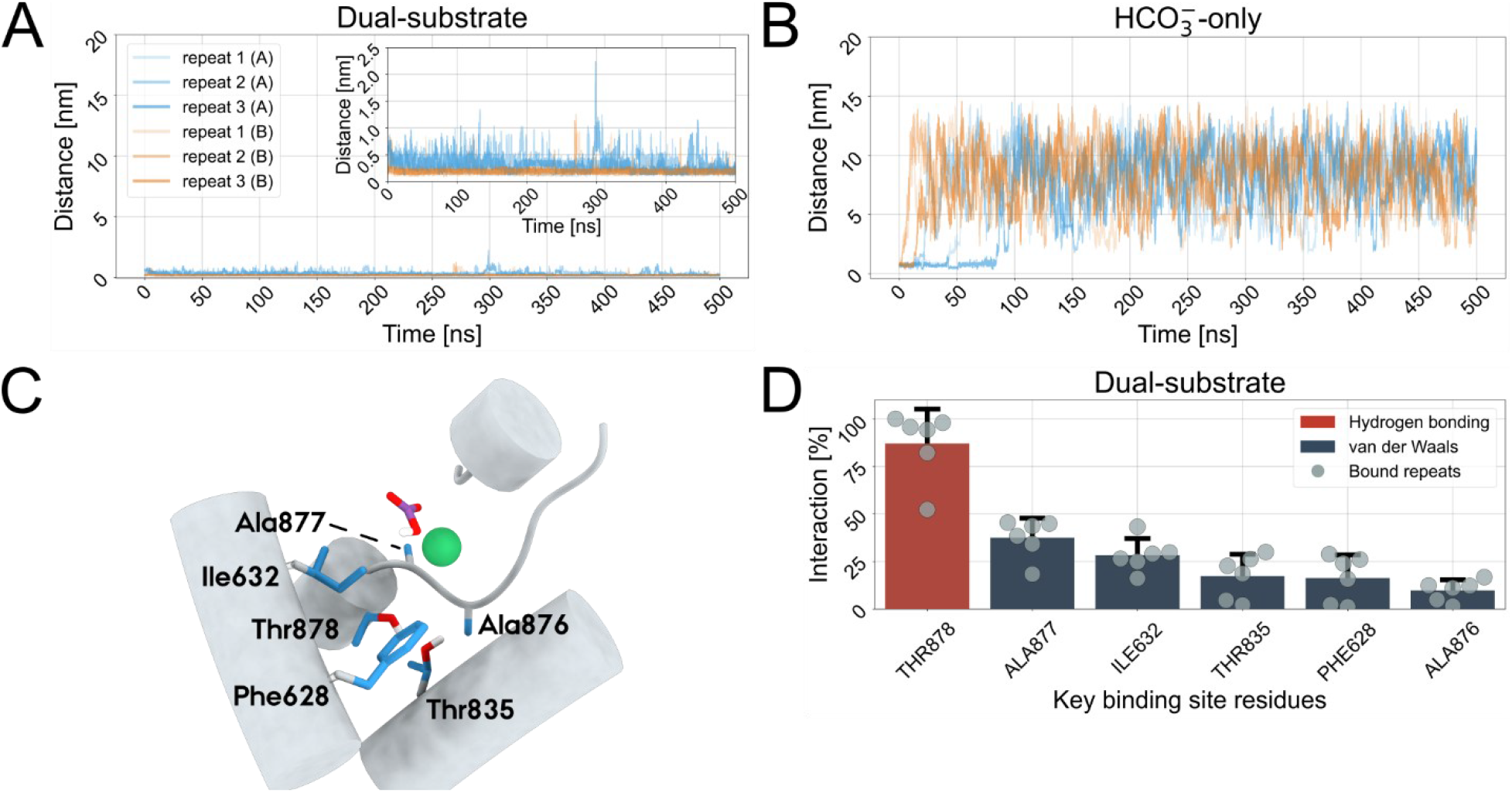
Stability and binding of HCO_3_^−^. (A) Plots of the distance between the binding site and HCO_3_^−^, in the dual-substrate system; HCO_3_^−^ is stable throughout the simulations for both chain A and B and in all repeats. (B) Distance plots for HCO_3_^−^ in the HCO_3_^−^-only system; HCO_3_^−^ dissociates from the binding site when Na^+^ is not present for both chain A and B, and in all repeats. The legend is the same in both (A) and (B), except for the inset in (A) showing the plot with a magnified distance-axis. (C) The key binding site residues identified to interact with HCO_3_^−^ for at least 5% of the total HCO_3_^−^-protein interactions over all simulations. The key binding site residues are highlighted in blue, HCO_3_^−^ in purple, and Na^+^ in green. (D) The interaction percentages for each key binding site residue and its interaction type. Data is included from all simulations where HCO_3_^−^ remains bound (i.e. all repeats in the dual-substrate systems), with data from each repeat shown individually as circles on the bar plot.

The key binding site residues of the dual-substrate system are Phe628, Ile632, Thr835, Ala876, Ala877, and Thr878 (Figure 2C). These are hydrophobic (phenylalanine, isoleucine, and alanine) and polar (threonine) residues. The main interaction residue in NBCn2 is Thr878, and it performs hydrogen bonding for 87% ± 18 of the total simulation time (Figure 2D). All other residues in NBCn2 participate through vdW interactions and serve to sterically stabilize HCO_3_^−^ in the binding site. HCO_3_^−^ also interacts with Na^+^ for the majority of the simulation time through ionic interactions (97% ± 2 of the total simulation time). Without Na^+^ (HCO_3_^−^-only), HCO_3_^−^ unbinds rapidly, meaning no key binding site residues can be identified in the absence of Na^+^. Losing the ionic interaction with Na^+^ destabilizes the HCO_3_^−^, and the residues in NBCn2 cannot compensate for the loss of Na^+^, and thus HCO_3_^−^ unbinds from the binding site.

### Na^+^ is affected by the presence of HCO3-in the binding site

The Na^+^ stability and binding were analyzed similarly to HCO_3_^−^ by performing MD simulations of two systems containing Na^+^ in the binding site (Table 1).

Na^+^ was stable throughout the simulations and across repeats in the presence of HCO_3_^−^ (dual-substrate system), similar to that of HCO_3_^−^ in the same system (Figure 3A). The distance value for Na^+^ is below 1.0 nm, which indicates that Na^+^ stays bound, irrespective of repeat and chain (Figure 3A zoom). Removing HCO_3_^−^ from the binding site (Na^+^-only system) resulted in Na^+^ unbinding in 50% of the repeats (two repeats from chain A and one repeat from chain B) (Figure 3B). This indicates that Na^+^ is able to stay bound in the binding site of NBCn2 without HCO_3_^−^ to a certain extent. However, the simulations do show that HCO_3_^−^ is able to affect the stability of Na^+^ binding. The residues in NBCn2 that interact with Na^+^ in the dual-substrate system are Thr569, Asp831, Thr835, Val875, Ala876, Ala877 and Thr878 (Figure 3C). These residues are hydrophobic (valine and alanine), polar (threonine), and negatively charged (aspartic acid). Removing the HCO_3_^−^ (Na^+^-only system) results in Na^+^ rapidly unbinding in half of the simulations, and for these repeats, no key binding site residues could be identified. For the remaining repeats, key binding site residues were identified, and they are primarily the same residues as in the dual-substrate system, however, some residues lost their interaction with Na^+^, namely Thr569 and Thr835 (Figure 3D). Some residues interact with both HCO_3_^−^ and Na^+^, and these are Thr835, Ala876, Ala877, and Thr878, where Thr878 was identified as the main interacting residue of HCO_3_^−^. The main interacting residue of Na^+^ in the dual-substrate system is Asp831, and it can perform ionic (100% ± 1 of the total simulation time) interactions with Na^+^. The remaining residues perform vdW interactions, which sterically stabilize Na^+^ in the binding site (Figure 3E). Asp831 is also the main interacting residue in the Na^+^-only system, and it interacts through ionic (51% ± 46 of the total simulation time) interactions. The reason for the large standard deviations stems from taking the mean of a total of six independent results, where three do not contribute to the calculation of the key binding site residues (Figure 3F). These results also reveal that Na^+^ does not interact with other residues in NBCn2 except when HCO_3_^−^ is present, hereby confirming that Na^+^ can be stable in the binding site on its own.

**Figure 3.**
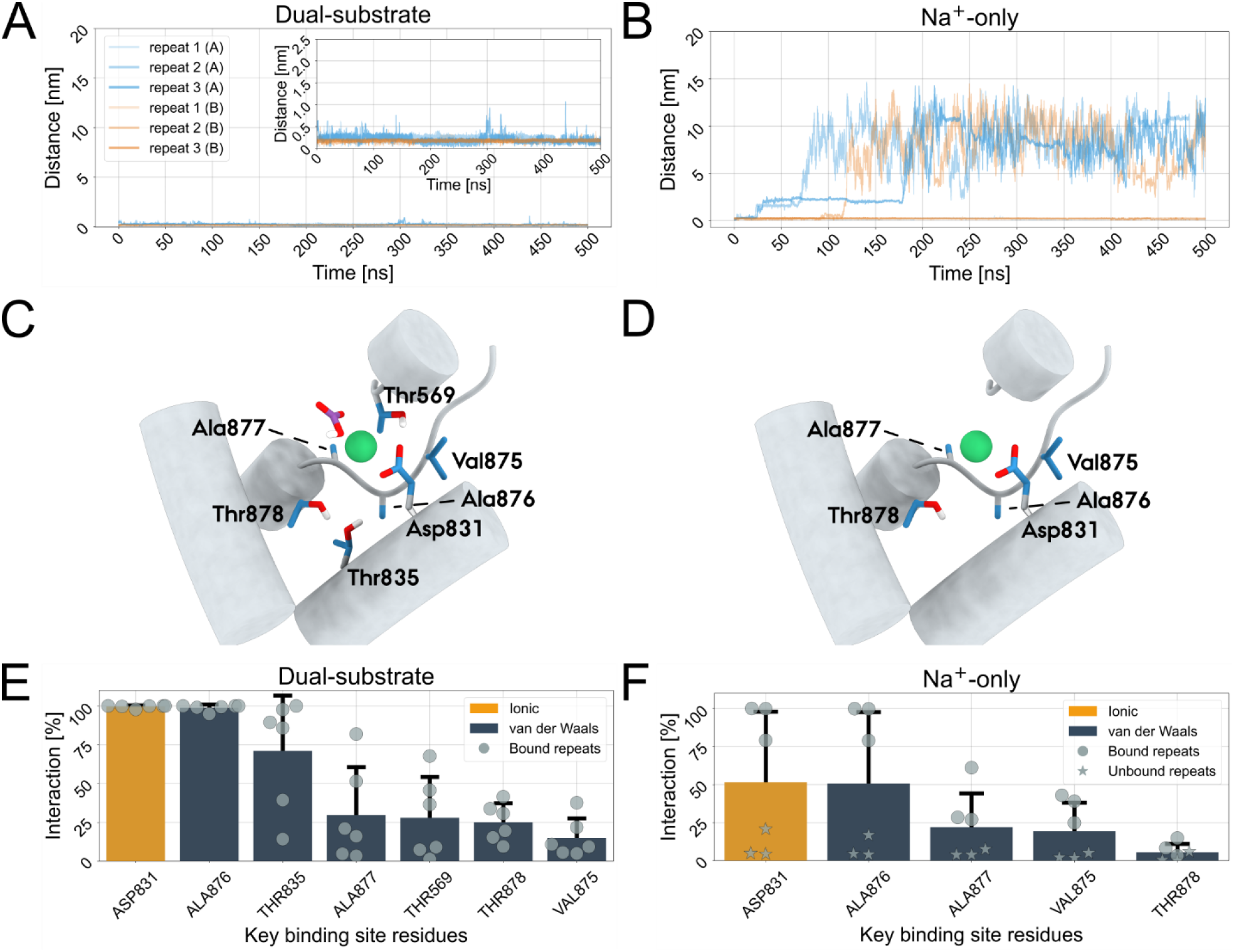
Stability and binding of Na^+^. (A) Distance for Na^+^ in the dual-substrate system (with respect to the binding site); Na^+^ is stable throughout the simulations for both chain A and B and in all repeats. (B) Distance for Na^+^ in the Na^+^-only system; Na^+^ dissociates from the binding site in half of the repeats in the absence of HCO_3_^−^. The legend is the same in both (A) and (B) except for the inset in (A), showing the plot with a magnified distance-axis. (C) The key binding site residues identified to interact with Na^+^ for at least 5% of the total Na^+^-protein interactions over all simulations in the dual-substrate system. (D) The key binding site residues identified to interact with Na^+^ for at least 5% of the total Na^+^-protein interactions over all simulations in the Na^+^-only system. The key binding site residues are highlighted in blue, HCO_3_^−^ in purple, and Na^+^ in green. (E) The interaction percentages for each key binding site residue and its interaction type in the dual-substrate system. The circles indicate the bound repeats. (F) The interaction percentages for each key binding site residue and its interaction type in the Na^+^-only system. The circles indicate that Na^+^ stayed bound in the binding site and the stars indicate that Na^+^ unbound from the binding site.

### Key binding site residues are conserved throughout the SLC4A family

The conservation of the key binding site residues that interacted with HCO_3_^−^ and Na^+^ were analyzed by performing multiple sequence alignment (MSA) of all ten members of the SLC4A family (Figure 4). In all instances, high conservation was found between the family members, except for the SLC4A11 protein, which is the least conserved SLC4A member^22^. SLC4A11 has a tyrosine in place of Thr878, which is the principal binding residue in the HCO_3_^−^ site in our simulations. Thus, SLC4A11 is not considered further in the present context. Almost all the residues found to interact with HCO_3_^−^ were conserved among the other family members apart from Ala877, which is a threonine in SLC4A1, and Ala876, which is a serine in SLC4A9. Both changes are from a hydrophobic residue to a polar residue. The residue numbering stems from the SLC4A10 protein. Residues that interacted with Na^+^ (apart from those that interacted with both Na^+^ and HCO_3_^−^ in the dual-substrate system) were found to be different, namely Thr569, Asp831, and Val875. Thr569 is a serine, and Asp831 is a glutamic acid in SLC4A1, A2, and A3. Val875 is a serine in SLC4A1, an alanine in SLC4A2, and a threonine in SLC4A3.

**Figure 4.**
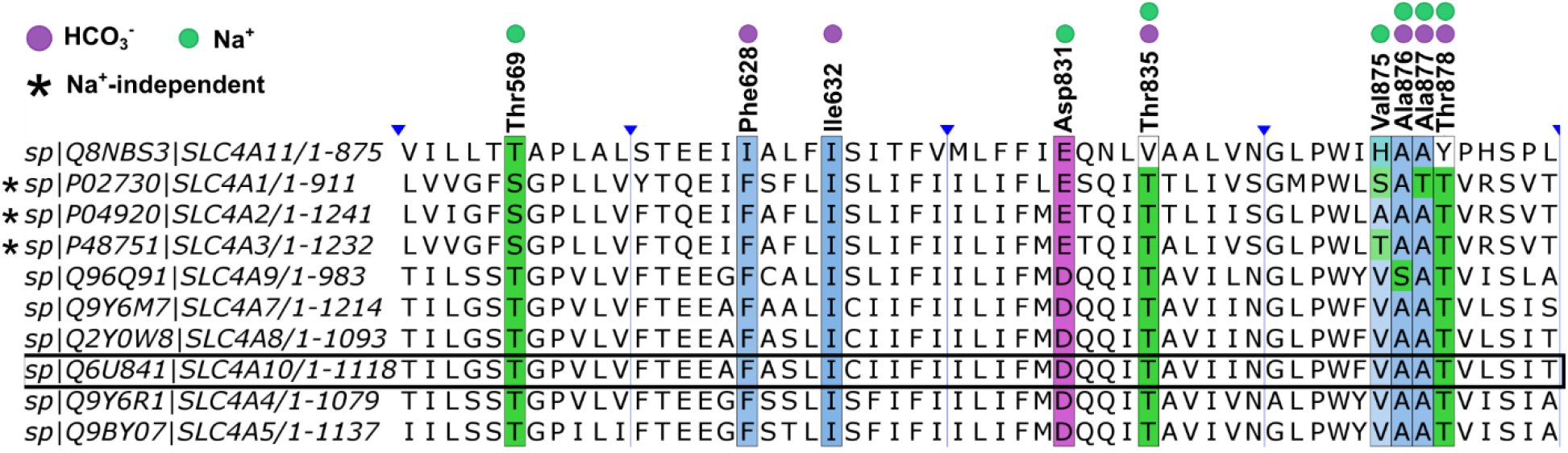
Conservation of key binding site residues. The multiple sequence alignment of the 10 members of the human *SLC4A* family. NBCn2 has been highlighted with a black rectangle, and the residue numbering stems from this protein. The residues have been colored using the Clustal coloring scheme according to their residue type (hydrophobic residues in blue, polar residues in green, and negatively charged residues in magenta), and the opacity is based on the degree to which the residue is conserved. The sequence alignment shows only the key binding site residues and five residues on either side of each key binding site residue; the blue arrows indicate parts of the sequences that have been hidden. The green circles indicate that the residue is a Na^+^ key binding site residue, the purple circles indicate that the residue is an HCO_3_^−^ key binding site residue, and the star indicates Na^+^-independent transporters.

### Na^+^-dependent HCO_3_^−^ transport data have been derived from pH measurements

Previous studies demonstrated that the transport of HCO_3_^−^ by NBCn2 is not only Na^+^-dependent but actually coupled to Na^+^ translocation in apparent symport with HCO_3_^−23^. This relationship is illustrated by the data from parallel *in vitro* measurements of intracellular [Na^+^] and pH with or without Na^+^ in the environment (Figure 5). The left side panel shows the [Na^+^]_i_ over time. After time in Na^+^ free medium, the estimated concentration is lower than normal levels in both cells with and without mouse Slc4a10 (lower case for mice protein due to naming convention). When Na^+^ is introduced into the medium, the [Na^+^]_i_ increases towards a normal physiological level within 30 seconds, but only in cells expressing Slc4a10, indicating inward Na^+^ transport by Slc4a10 protein in the presence of HCO_3_^−^. The right side panel shows the corresponding pH_i_ changes by the same experimental protocol. The initial pH_i_ in the absence of Na^+^ is low in both cells with and without Slc4a10 protein. When Na^+^ is introduced, the pHi increases towards normal physiological levels only in cells expressing Slc4a10, indicating inward HCO_3_^−^ transport by Slc4a10 protein in the presence of Na^+^. The rate of pH_i_ changes seems to be slow compared to the [Na^+^]_I_ increase because of the pH buffering capacity of the cells. The calculated rates of Na+ and HCO_3_^−^ influx (derived from the apparent H^+^ efflux) are comparable.

**Figure 5.**
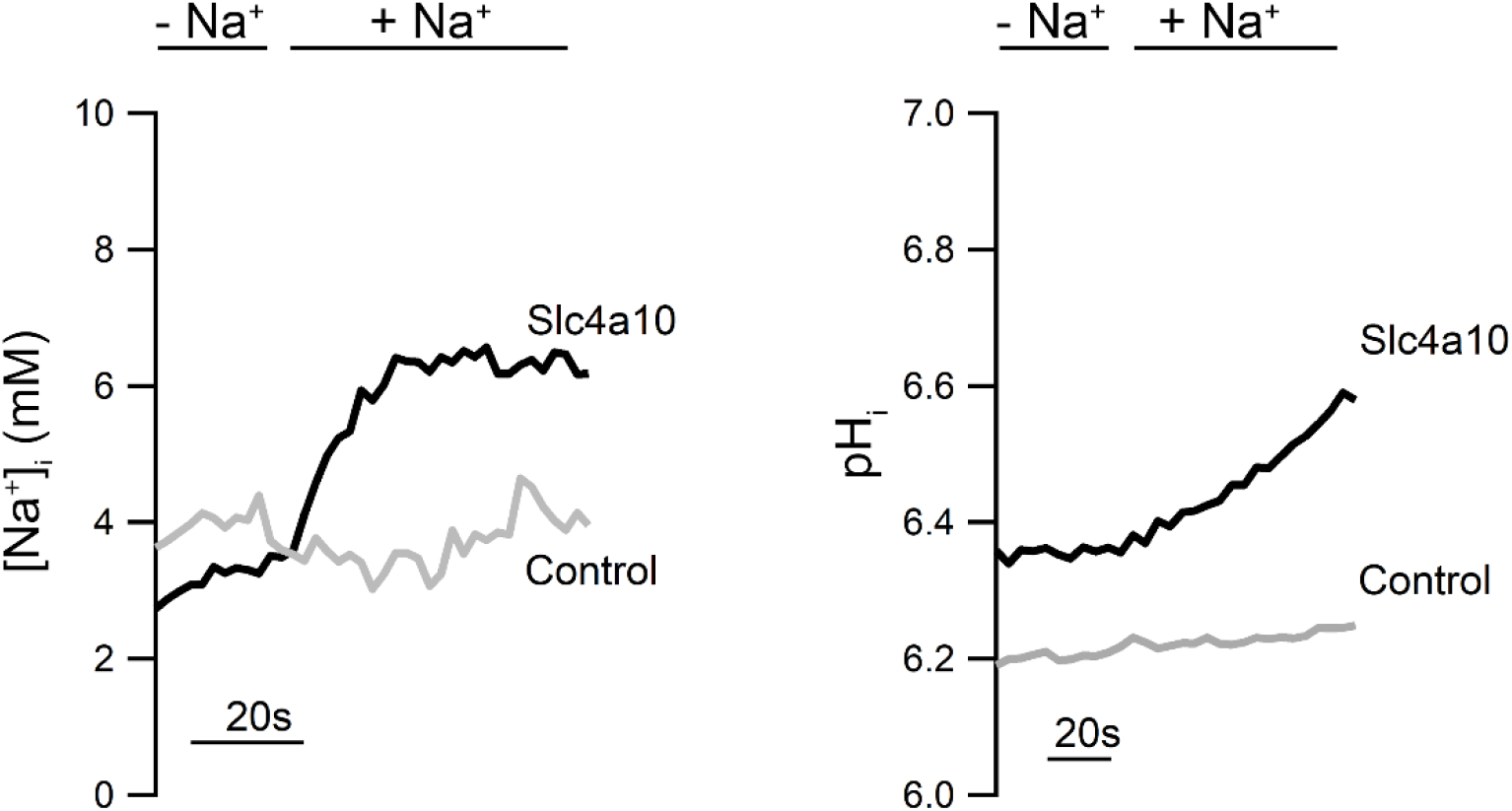
Na^+^-dependent HCO_3_ ^−^ transport. The left panel shows the time course of intracellular [Na^+^] ([Na^+^]_i_) changes in control and the mouse homolog of NBCn2/Slc4a10-transfected NIH-3T3 cells were superfused with Na^+^-free (-Na^+^) solution before 145 mM Na^+^ was reintroduced (+Na^+^) to the bath solution as indicated. The right panel shows the corresponding changes in intracellular pH (pH_i_) during the same procedure. 5% CO_2_ and 25 mM HCO_3_^−^ were present throughout the procedures. Note that the apparently slower pH_i_ response than [Na^+^]_i_ response is a consequence of the intracellular pH buffering.

## DISCUSSION

The MD simulations demonstrated the necessity of Na^+^ for HCO_3_^−^ stability, as HCO_3_^−^ unbinds rapidly from the binding site in the absence of Na^+^. Probing the importance of HCO_3_^−^ for Na^+^ binding revealed that the absence of HCO_3_^−^ from the binding site does not necessarily lead to Na^+^ unbinding, but the presence of HCO_3_^−^ still stabilizes the binding. We find that Na^+^ is located deeper in the binding pocket of NBCn2 than HCO_3_^−^, which could further explain the increased stability of Na^+^ and differentiation between the two ions in terms of their dependence on one another for stability. We posit that this allows Na^+^ to stay bound by the key binding site residues despite losing the ionic interaction with HCO_3_^−^, though with reduced stability.

### Structure prediction and MD simulations

The generated model for NBCn2 was based on the available structural data for the SLC4A family. All members of the SLC4A family are functional homodimers, and NBCn2 was modeled accordingly^5^. Due to the high sequence homology of the SLC4A family, the proteins can be expected to have a similar overall fold^24^. Accordingly, evidence in support of the SLC4A10 structure was obtained using AlphaFold2, which resulted in an overall pLDDT score of 92 for the NBCn2 homodimer. Thus, we are confident in our structural model as a starting point for simulations.

### Key binding site residues correlate with known interactions

The Asp831, Thr835, and Thr878 have been shown to be important in coordinating Na^+^ in another study of SLC4A10 and in SLC4A8^25^ further cementing the importance of Na^+^ presence. The serine on position 569 (SLC4A10 numbering) and the glutamic acid on position 831 (SLC4A10 numbering) in SLC4A1 has been shown to be important for transport^26-28^. These residues are notably different than the ones observed for Na^+-^dependent transport, where they are threonine and aspartic acid, respectively. Another interesting residue found in SLC4A1 that is crucial for transport is an arginine in position 880 (SLC4A10 numbering). This residue is conserved for the three Na^+^-independent transporters, SLC4A1, A2, and A3. In SLC4A4, A5, and A9, it is an isoleucine, while in SLC4A7, A8, and A10, it is a leucine. This further highlights the chemical nuances that distinguish the SLC4A family of transporters and divides them into their subfamilies. Recent work on the SLC4A9 protein has shed some light on important residues for transport activity. This study found that when mutating Asp831, Thr569, and Thr878 (SLC4A10 numbering), HCO_3_^−^ transport was reduced^29^. These residues are highly conserved in Na^+^-dependent transporters of the SLC4A family, and they were also revealed to be critical for coordination and binding in NBCn2 by our MD simulations. It is noteworthy that the residues interacting with HCO_3_^−^ are highly conserved throughout the *SLC4A* family, which corresponds to universal transport of HCO_3_^−^. High sequence conservation was observed throughout the family members except for the SLC4A11 protein, which is the only member of the *SLC4A* family that does not transport HCO_3_^−^. The SLC4A11 substrate is still under investigation^30-32^.

### Na^+^-independence in SLC4A1 to 3

Across the SLC4A family, the Na^+^ key binding site residues were less conserved than the HCO_3_^−^ residues, and particularly they differed from SLC4A10 in SLC4A1, A2, and A3, which are known to be Na^+^-independent HCO_3_^−^ transporters. This may explain why Na^+^ is not bound to the binding sites of these proteins. The change of Asp831 (SLC4A10 numbering) to a glutamic acid in SLC4A1, A2, and A3 does not alter the charge of the side chain, but perhaps changes the volume and coordination in the binding site, which may hinder Na^+^ binding. The change of Thr569 to a serine keeps the sidechain as a polar residue, but again, perhaps this alters the coordination environment of the binding site. The change of Val875 to a serine, alanine, or threonine, in SLC4A1, A2, and A3, respectively, changed the sidechain from a hydrophobic residue to a polar residue in SLC4A1 and SLC4A3, while keeping it hydrophobic in SLC4A2.

### Substrate stoichiometry

The shown example experiment of intracellular pH and Na^+^ changes in cells expressing Slc4a10 is summarized in our previous report^23^ demonstrates the absolute dependence of HCO_3_^−^ import into cells on extracellular Na^+^, and that the transport of the two ions is in the same direction. In that study, the requirement of Na^+^ import for HCO_3_^−^ is also shown^23^. This finding correlates with the MD simulation results here, showing that HCO_3_^−^ is only stable and thus able to be transported by NBCn2 when Na^+^ is present in the binding site to stabilize it. The reported apparent stoichiometry for the mouse Slc4a10-mediated cotransport was 1Na^+^:2HCO_3_^−^, which is experimentally indistinguishable from 1Na^+^:1CO_3_^2-^. The electroneutrality was maintained by obligate antiport with Cl^−^ and, thus, behaves as a Na^+^ dependent Cl^−^ /HCO_3_^−^ exchanger in rodents, as reported by others^25,33,34^. Conversely, the human SLC4A10 seems to operate independently of an apparent Cl^−^/Cl^−^ self exchange^3^. Such interspecies differences in transport substrates and stoichiometries are not uncommon, and have been reported for e.g., solute carrier families, especially the SLC26 family of more promiscuous anion transporters^35^. Moreover, an elegant experimental study demonstrated that the human electrogenic Na^+^:HCO_3_^−^ transporter SLC4A4 (NBCe1) translocates CO_3_^2-^ along with 1 Na^+^, while the anion exchanger SLC4A1 (AE1) indeed exchanges Cl^−^ for HCO_3_^−36^. It has not been experimentally tested whether SLC4A10 translocates HCO_3_^−^ or CO_3_^2-^. Thus, the transporters of this family might prefer different ionic species as a background for the differences in apparent stoichiometry. It is, thus, imperative to uncover the ion species preference of SLC4A10 in future experimental as well as computational analyses.

### Limitations

We should note that we cannot discriminate between CO_3_^2-^ and HCO_3_^−^ in this analysis, and that we do not consider counterion involvement. That is: we deliberately chose an in silico experimental design that would be relevant for both 1Na^+^:1HCO_3_^−^ cotransport and 1Na^+^:1CO_3_^2-^/1Cl^−^ exchange transport modes. In this study, we describe a simple membrane in which the protein is embedded. Biological membranes consist of a large variety of different lipids depending on the cell type and environment. The lipid composition of the membrane is known to affect membrane proteins and could therefore be of potential relevance^37-40^. Due to sampling constraints of the simulations, we did not observe any conformational changes of NBCn2 towards the inward-open state. We only performed simulations with protein sidechain charges assigned by PROPKA, and it would be interesting to study how different protonation states would impact the substrate when bound in the binding site. To this end, QM/MM^41,42^ and constant pH^43-46^ simulations would be of high interest to understand how hydrogen exchange/bonding between the HCO_3_^−^ and protein sidechains occurs.

### Conclusion and perspective

Based on simulation data and pH experiments that we present here, we propose a model for the substrate binding of HCO_3_^−^ and Na^+^ that leads to their transport by NBCn2 (See Figure 6). NBCn2 starts in its apo-state and outward-open conformation and can bind Na^+^ and HCO_3_^−^. Na^+^ is bound first by NBCn2 (Figure 6 step II) and then HCO_3_^−^ is bound (Figure 6 step III). It is necessary for Na^+^ to bind first, since it is located deeper in the binding cavity than HCO_3_^−^ (Figure 6 step III zoom in). NBCn2 undergoes a conformational change from its outward-open state to its inward-open state, hereby translocating the substrate (Figure 6 step IV). The substrate is released in the intracellular cytoplasm, and NBCn2 is in its apo-state in the inward-open conformation (Figure 6 step V). Finally, NBCn2 undergoes a conformational change back to its apo-state. The proposed sequential binding model could potentially be how all Na^+^-dependent transporters of the SLC4A family bind their substrate. A potential way of confirming the proposed sequential binding mechanism would be to mutate each binding site residue one at a time, to observe if the binding stability would be impacted by a single point mutation. Either knocking out the Na^+^ or HCO_3_^−^ specific residues in both computational models and *in vitro* could help reveal how the various residues aid in the binding and coordination of the substrate. The exact pathway through which the substrate is translocated through the protein is still unknown and is required to fully map the transport mechanism of NBCn2. To obtain this information, enhanced sampling simulations would prove beneficial, as it is possible to apply an external bias to the protein, thereby driving it between its conformational states. This would also aid in the determination of the exact substrate for NBCn2 transport.

**Figure 6.**
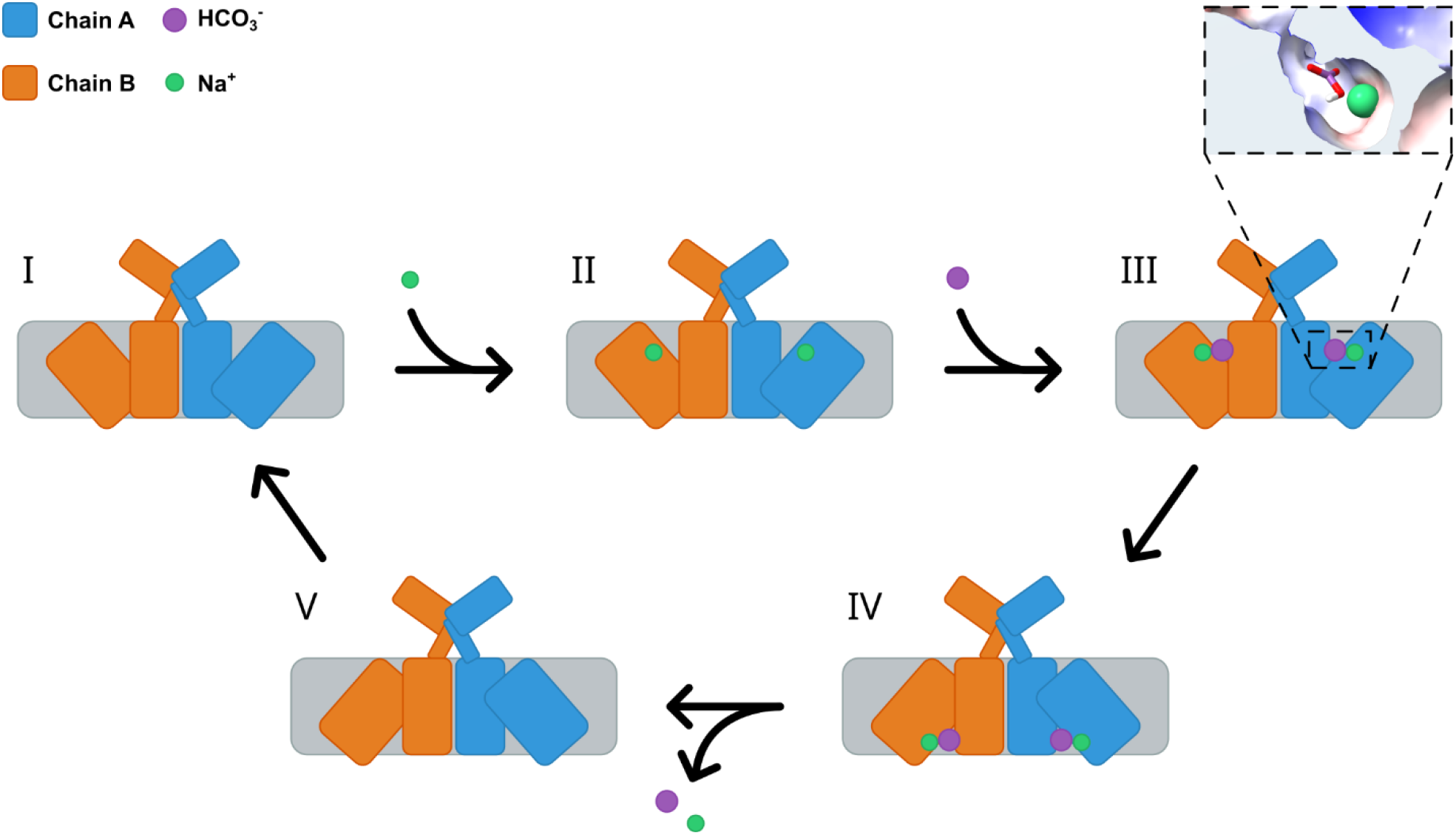
Sequential binding of substrates to NBCn2. Proposed sequential binding mechanism of NBCn2. I) The apo-state of NBCn2 in outward-open conformation. II) Binding of Na^+^ in the binding site of both chains. III) Subsequent binding of HCO_3_^−^ in the binding site of both chains. The insert shows how Na^+^ and HCO_3_^−^ fit in the binding site cavity. Na^+^ is found to bind deeper in the binding site and hence needs to bind before HCO_3_^−^ to stabilize it. IV) Conformational change of the protein through an elevator-type mechanism, where the ions are translocated through the protein chains. V) Release of Na^+^ and HCO_3_^−^ into the cell, which leads to an apo-state in the inward-open conformation. Final recovery back to apo-state in the outward-open conformation. Chain A in blue, chain B in orange, Na^+^ in green, and HCO_3_^−^ in purple.

## RESOURCE AVAILABILITY

### Lead contact

Requests for further information and resources should be directed to and will be fulfilled by the lead contact, Anna Louise Duncan

### Materials availability

This study did not generate new unique reagents.

### Data and code availability

Simulation coordinate data have been deposited at Zenodo and are public as of the date of publication, the DOI is listed in the key resources table. This paper does not report original code. Any additional information required to reanalyze the data reported in this paper is available from the lead contact upon request.

## ACKNOWLEDGMENTS

L.M.D. acknowledges support from Aarhus University Research Foundation (AUFF Nova; AUFF-E-2022-9-24). A.D.S. is supported by grants from Danmarks Frie Forskningsfond (grant no. 0136-00148B) and the Novo Nordisk Foundation (NNF20OC0065431). L.M.D., A.D.S., B.S., and A.L.D. had access to computational resources at the Grendel cluster of the Centre for Scientific Computing Aarhus, and the Resource for Biomolecular Simulations (ROBUST; supported by the Novo Nordisk Foundation; NNF18OC0032608, NNF24OC0087976). J.P. and H.H.D. were supported financially by the Danish Medical Research Council (10-080511), the Lundbeck Foundation (R32-A2838), and Aarhus University Research Foundation (F-2009-SUN-1-43).

## AUTHOR CONTRIBUTIONS

Conceptualization, L.M.D., J.P, B.S. and A.L.D.; methodology, L.M.D., A.D.S., H.H.D, and J.P.; investigation, L.M.D., A.D.S, H.H.D, and J.P.; writing – original draft, L.M.D., J.P., and A.L.D; writing – review & editing, L.M.D., A.D.S., B.S., H.H.D., A.L.D., and J.P.; funding acquisition, B.S. and J.P.; resources, A.L.D., H.H.D., and J.P.; supervision, B.S., A.L.D, and J.P.

## DECLARATION OF GENERATIVE AI AND AI-ASSISTED TECHNOLOGIES

During the preparation of this work, the author(s) used ChatGPT to check for grammatical errors in the writing. After using this tool or service, the author(s) reviewed and edited the content as needed and take(s) full responsibility for the content of the publication.

## METHOD DETAILS

### Modelling

#### Sequence truncation

The sequence used for structure prediction was based on the human *SLC4A10* protein using UniProt ID: Q6U841^47^. The sequence was 1118 amino acids long and was truncated in both the C- and N-terminal. The truncation was based on pairwise sequence alignment using EMBOSS NEEDLE^48^ with the sequence from the Ndcbe structure (SLC4A8 protein) with PDB ID: 7RTM^14^. This resulted in a sequence of 554 amino acids, meaning the N-terminal was truncated by 481 residues and the C-terminal was truncated by 83 residues. NBCn2 is functionally a homodimer, which results in a total length of 1108 amino acid residues.

#### Structure prediction

Colabfold^21,49,50^ (AlphaFold2) was used to generate the starting structure of the NBCn2 homodimer. Multiple templates were used as input for the structure prediction. Only structures from the *SLC4A* family were used, and only if they were determined to be in the outward-open conformation. The structures used for structure prediction can be found in Table 2. The max-msa was set to 32:64 to reduce the model’s knowledge about the MSA, thereby forcing it to rely more on the provided templates. The model-type was set to alphafold2_multimer_v3, num-recycle to 10, and num-relax to 5 using Amber. The top-scoring (rank 1) structure was chosen as the starting structure.

**Table 2.** Structural templates.

| Protein name | Gene name | PDB ID(s) | Publication |
| --- | --- | --- | --- |
| AE1 | SLC4A1 | 7TYA, 7TY4, 7TY6, 7TY7, 7TY8, 8T6U, 8T6V | Capper et al.<sup>8</sup> |
| AE1 | SLC4A1 | 7UZ3, 7V07, 7V19, 8CRQ, 8CRR, 8CT3 | Vallese et al.<sup>9</sup> |
| AE1 | SLC4A1 | 7TVZ, 7TW0, 7TW1, 7TW3, 7TW5, 7TW6 | Xia et al.<sup>10</sup> |
| AE2 | SLC4A2 | 8GV8, 8GVC, 8GVF | Zhang et al.<sup>11</sup> |
| AE3 | SLC4A3 | 8Y85, 8Y86, 8Y8K | Jian et al.<sup>12</sup> |
| Nbce1 | SLC4A4 | 6CAA | Huynh et al.<sup>13</sup> |
| Ndcbe | SLC4A8 | 7RTM | Wang et al.<sup>14</sup> |
| Btr1 | SLC4A11 | 7X1I, 7X1J | Lu et al.<sup>15</sup> |

#### System preparation

The protein was prepared using the Protein Preparation Wizard^51^ in Schrodinger’s Maestro v2021.4^52^. Protein protonation states were assigned using PROPKA^53-55^ at a pH of 7.4. The three different system setups (dual-substrate, HCO_3_^−^-only, and Na^+^-only) were prepared by taking the coordinate values for HCO_3_^−^ and Na^+^ from the Ndcbe (SLC4A8) structure with PDB ID: 7RTM^14^. This was done by superimposing the structure of Ndcbe on our starting structure and copying the coordinates for HCO_3_^−^ and Na^+^ into our starting structure. For the HCO_3_^−^-only system, Na^+^ was deleted, and for the Na^+^-only system, HCO_3_^−^ was deleted.

### Molecular dynamics simulations

#### System setup

The CHARMM-GUI Membrane Builder was used to generate the topology files for the MD simulations^56-58^, which consisted of the NBCn2 homodimer, a membrane of 56% POPC and 44% cholesterol, and solvated with TIP3 water. A salt concentration of 150 mM NaCl was added to the solvent, and 25 mM HCO_3_^−^ was added to the solvent, and the system was neutralized with NaCl. The setup was done automatically using the CharmmGuiAuto.py script^59^. All simulations were performed using the Amber SB19 Force field^58,60^ in GROMACS 2024.3^61,62^.

The systems were minimized using the steepest descent algorithm for 5000 steps. Afterward, six rounds of equilibration were performed. The first equilibration was performed in the NVT ensemble using a timestep of 1fs and a total simulation time of 0.125 ns using the Berendsen thermostat^63^. The second equilibration was performed in the NVT ensemble using a timestep of 1fs and a total simulation time of 0.125 ns using the Berendsen thermostat. The third equilibration was performed in the NPT ensemble using a timestep of 1fs and a total simulation time of 0.125 ns using the Berendsen barostat and thermostat. The fourth equilibration was performed in the NPT ensemble using a timestep of 2fs and a total simulation time of 0.5 ns using the Berendsen barostat^63^ and thermostat. The fifth equilibration was performed in the NPT ensemble using a timestep of 2 fs and a total simulation time of 0.5 ns using the Berendsen barostat and thermostat. The sixth equilibration was performed in the NPT ensemble using a timestep of 2 fs and a total simulation time of 0.5 ns using the Berendsen barostat and thermostat. This was done to gradually decrease the position restraints on the protein backbone and side chains and the lipids. The values were lowered from 4000 kJ/mol to 50 kJ/mol, 2000 kJ/mol to 0 kJ/mol, and 1000 kJ/mol to 0 kJ/mol, respectively. The production run was performed in the NPT ensemble using the Parinello-Rahman barostat^64^ and the v-rescale thermostat^65^. The temperature was set to 310 K, τ_t_ was 1.0 ps, the pressure was set to 1 bar, and τ_p_ was 5.0 ps, the compressibility was 4.5*10^−5^ bar^−1^. The PME type was used for the long-range electrostatic interactions, and for the short, the cutoff was set to 1.2 nm^66,67^. Van der Waals interactions were considered up to the cutoff of 1.2 nm using the force-switch modifier between 1.0 – 1.2 nm. The LINCS algorithm^68^ was used to constrain the hydrogen bonds. All simulations were produced with three replicas using a 2 fs timestep for 500 ns, yielding in total of 1.5 µs for each system, and a total of 4.5 µs for the simulations presented in this paper.

#### Analysis

The plots were generated using the matplotlib package^69^ and the seaborn package^70^ in Python. Structures were visualized using ChimeraX^71^. Protein – ion contact fingerprints and ion – ion contact fingerprints were generated using ProLIF^72^ and MDAnanlysis^73^. GROMACS analysis tool gmx pairdist was used to calculate the distance values for Na^+^ and HCO_3_^−^ with respect to the binding site^61^. The binding site was defined as all key binding site residues for both Na^+^ and HCO_3_^−^, which were Thr569, Phe628, Ile632, Asp831, Thr835, Val875, Ala876, Ala877, and Thr878. The distance was calculated between the center of mass of the nine binding site residues’ Cα-atoms and the center of mass of either Na^+^ or HCO_3_^−^. The two chains of NBCn2 are structurally identical and each binds both HCO_3_^−^ and Na^+^, and thus they will be accounted for as independent replicas in the results section but highlighted separately.

### Bioinformatics

To determine the conservation of the key binding site residues, multiple sequence alignment was performed using the Clustal Omega tool^48^. Sequences from all *SLC4A* family members were used (SLC4A1-5 and SLC4A7-11). The key residues were highlighted and colored based on their amino acid type and their degree of conservation using Jalview^74^.

### Experimental methods

#### Cell Culture

FRTs containing NIH-3T3 mouse fibroblasts (Invitrogen) were grown in Dulbecco’s modified Eagle’s medium with glutamax™ supplemented with 10% donor bovine serum. Cells were stably transfected with a single copy of a mouse *slc4a10* (AB033759) or control cDNA construct into a predefined locus using the Flp-InTM system (Invitrogen). Transfection and protein expression were previously validated by PCR and sequencing, surface biotinylation, Western blotting, and immunocytochemistry^23^.

#### Dynamic measurements of intracellular pH and [Na+]

Cells were grown to approximately 50% confluence on glass coverslips. For pH_i_ measurements, cells were loaded with 2 μM BCECF-AM in a HEPES-buffered salt for 15 min. For intracellular [Na^+^] measurements, cells were loaded in 10 μM CoroNa Green sodium indicator for 30 min in HBS. All incubations were performed in a dark chamber heated to 37 °C. Fluorophores were purchased from Invitrogen. The cells were then mounted in a closed perfusion chamber (358-μl RC-21BR or 36-μl RC-20; Warner Instruments) and perfused with a linear flow rate of 0.8 mm/s at 37 °C. For pH_i_ measurements, the dye excitation periods of 20 ms were alternated between 495-nm and 440-nm light from a monochromator (Till Photonics). The light emission at 510–535 nm was recorded by a 12-bit cooled monochrome CCD camera (QImaging, Retiga EXi). QED InVivo imaging software (Media Cybernetics) was used to control wavelength, light exposure time (20 ms), frequency (one image pair every 4 s), and 4 × 4 binning (to 348 × 260 pixel images), as well as the data collection from regions of interest (one/individual cell). The mean values for cells from one coverslip represent n = 1. The excitation fluorescence ratio derived from the BCECF measurements was calibrated to intracellular pH_i_^23^. For the single wavelength [Na^+^]_i_ dye CoroNa Green, the excitation/emission wavelengths were 492/516 nm. CoroNa fluorescence was calibrated to [Na^+^]_i_ values by stepwise changes in extracellular [Na^+^] in the presence of the ionophore monensin, 10 μM.

## QUANTIFICATION AND STATISTICAL ANALYSIS

The NumPy Python package was used to carry out the statistical analysis^75^. It was used to determine the mean and standard deviation of the residue contacts to the ions. The three repeats and two monomeric chains were averaged, and the standard deviation was found based on this mean. This was plotted using the built-in function in seaborn^70^ as error bars in the bar plots. Data acquired using QED InVivo imaging (Media Cybernetics) was exported to IGOR Pro (WaveMetrics) for quantitation and analysis.

